# Modular Engineering of *Escherichia coli* for Enhanced Nickel Uptake, Survival, Biomineralization, and Hydrogen-Supported Bioremediation

**DOI:** 10.1101/2025.04.13.648562

**Authors:** Yi Shi, Ruiwen Ma, Lin Qi, Xujie Tan, Liye Chen, Yuhan Wang, Kexin Huang, Ziming Suo, Yuhan Li, Hongcheng Chen, Zhengchan Chen, Xueting Chen, Wangli Li, Kexin Zhen

**Affiliations:** Fudan University

## Abstract

Nickel contamination poses increasing environmental and industrial challenges, demanding effective and sustainable remediation strategies. Here, we present a modular synthetic biology approach to engineer *Escherichia coli* for efficient nickel enrichment, resilience in toxic environments, intracellular biomineralization, and hydrogen-based co-culture stability. Our system comprises four functional modules: (1) a **nickel enrichment module**, in which the fusion protein NixA-F1v outperformed existing uptake systems, and a mutant repressor RcnR^C35L^ enhanced retention by suppressing nickel efflux; (2) a **survival module** where heterologous expression of *Helicobacter pylori* Hpn conferred tolerance to high nickel concentrations, and YejM increased phage resistance; (3) a **nickel microparticle module** enabling intracellular bioconversion of nickel into detectable microparticles even without engineered uptake proteins; and (4) a **hydrogen supply module** supporting co-culture stability through *E. coli*-cyanobacteria adhesion for sustained hydrogen production. These modules were experimentally validated through comparative uptake assays, survival profiling, and microscopy-based particle detection. Our findings demonstrate a customizable and integrative microbial platform for metal bioremediation and biosensing, with potential applications in environmental engineering and industrial waste treatment.

## Introduction

Nickel (Ni) is extensively utilized in various industrial applications. However, significant amounts of dissolved nickel ions are discharged into the environment via municipal wastewater, landfill leachates, and mining runoff, posing substantial risks to ecosystems due to the lack of efficient recycling mechanisms. In soil environments, excessive nickel concentrations can hinder plant growth and reduce agricultural yields, while in aquatic systems, nickel toxicity impacts various organisms across trophic levels^[1][2]^. Notably, the most affected species include gastropods, cladocerans, and vascular plants.^[3]^ Thus, addressing the recovery of free nickel ions is crucial to mitigating these environmental threats.

Traditional recovery methods primarily focus on extracting nickel from discarded products such as stainless steel and batteries, using techniques such as smelting, refining, or chemical leaching of scrap metal. While these processes contribute to nickel recovery and mitigate its environmental impact, they are often energy-intensive, costly, and generate secondary waste. As a result, recycled nickel accounts for only a small fraction of the global supply, with mining operations—motivated by stronger economic incentives—continuing to exacerbate environmental degradation.^[4]^

Biological recovery methods present an affordable and sustainable alternative for treating heavy metal pollution, including nickel. However, several challenges must be addressed before these methods can be broadly implemented. Most microorganisms lack the capacity to selectively and actively accumulate substantial quantities of specific metal ions. High external concentrations of nickel also pose significant toxicity risks to microbial survival, while intracellular accumulation further complicates the recovery process. Moreover, effective collecting techniques must be developed to prevent the reintroduction of nickel into the environment following microbial death, and ideally, to convert the metal into a usable industrial form. These issues present considerable obstacles to the development of bio-recovery technologies.

Despite these challenges, solutions have been proposed. Several nickel-specific active transporters have been identified, and methods to inhibit the intrinsic nickel efflux mechanisms in Escherichia coli have been developed. ^[5][6][7]^The expression of metallothioneins has been shown to enhance microbial tolerance to high concentrations of heavy metals.^[8]^ Studies on hydrogenases further offer potential pathways for reducing metal ions, including nickel.^[9]^ In this study, we apply a synthetic biology approach to integrate these capabilities, thus facilitating efficient recovery and reuse of nickel contamination.

## RESULTS

### Engineering of a High-Efficiency Nickel Uptake System

To enhance nickel accumulation in *Escherichia coli*, we engineered and tested several nickel uptake constructs, including the native nik operon, nik operon coupled with a ribozyme regulation element (nik-ribozyme), the *Helicobacter pylori* nickel transporter NixA, a chimeric variant F1v-NixA, and a reverse fusion NixA-F1v. Nickel uptake was quantified in engineered strains cultured in media supplemented with 1 mM NiSO_4_. Among all variants, NixA-F1vdemonstrated the highest nickel uptake efficiency, significantly outperforming both the canonical nik operon and NixA alone (Figure 1). This result suggests that the C-terminal fusion of the F1v peptide to NixA enhances membrane insertion or nickel translocation activity, making NixA-F1v the optimal candidate for downstream applications.

**Figure 1:**
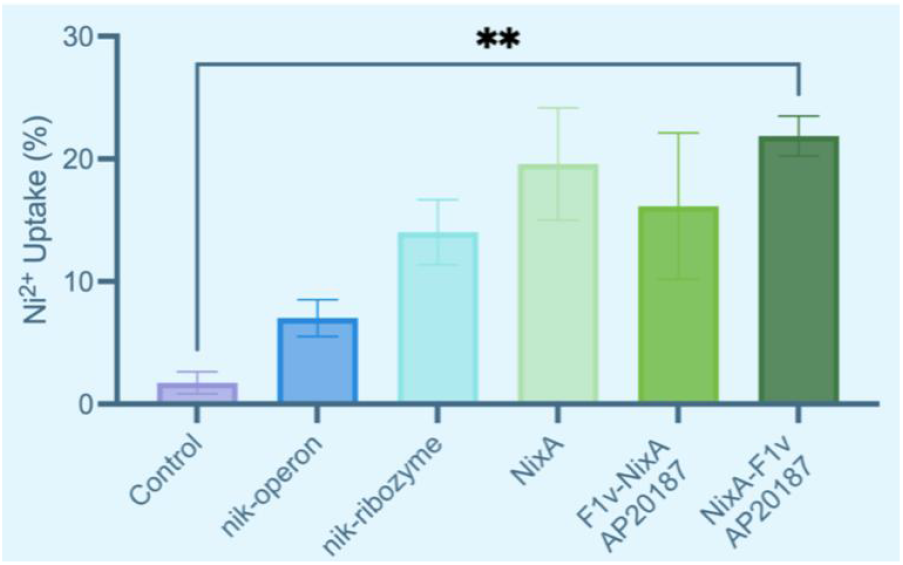
Comparison of Ni^2+^ Uptake Efficiency by Different *E. coli* in 30 mg/L Ni^2+^. The graph shows the percentage of Ni^2+^ absorbed by *E. coli* expressing different constructs after 5 hours of growth in a medium containing 30 mg/L Ni^2+^ (*E. coli* strain: BL21 DE3, induced with 1 mM IPTG). Ni^2+^ uptake was calculated based on the difference between initial and final concentrations in the supernatant, divided by 30 mg/L. The optical density (OD_600_) of the initial bacterial suspension was adjusted to 0.5. Culture for 5 hours, at 37°C with a rotating speed at 220 rpm. Regarding NixA-F1v and F1v-NixA, AP20187 is a synthetic dimerizer that can be used to induce homodimerization of F1v domain. Three biological replicates were performed for each condition, and error bars represent the standard errors of the means (SEM) of these replicates. ANOVA test shows that all constructs increase Ni^2+^ uptake significantly compared to the control. Bacteria expressing NixA-F1v exhibit the highest Ni^2+^ uptake efficiency (p = 0.0052, Dunnett’s post-test).

### Inhibition of Nickel Efflux via RcnR^C35L^

To further improve intracellular nickel retention, we introduced a point mutation in rcnR, generating the C35L variant, which was hypothesized to reduce efflux activity by impairing the repressor’s ability to induce rcnA. Engineered strains expressing rcn^RC35L^exhibited markedly elevated intracellular nickel levels compared to those expressing wild-type rcnR or harboring the empty vector control (Figure 2). This result supports the role of rcnR^C35L^ in effectively suppressing efflux pump activity, thereby enhancing nickel accumulation when used in conjunction with the NixA-F1v uptake module.

**Figure 2:**
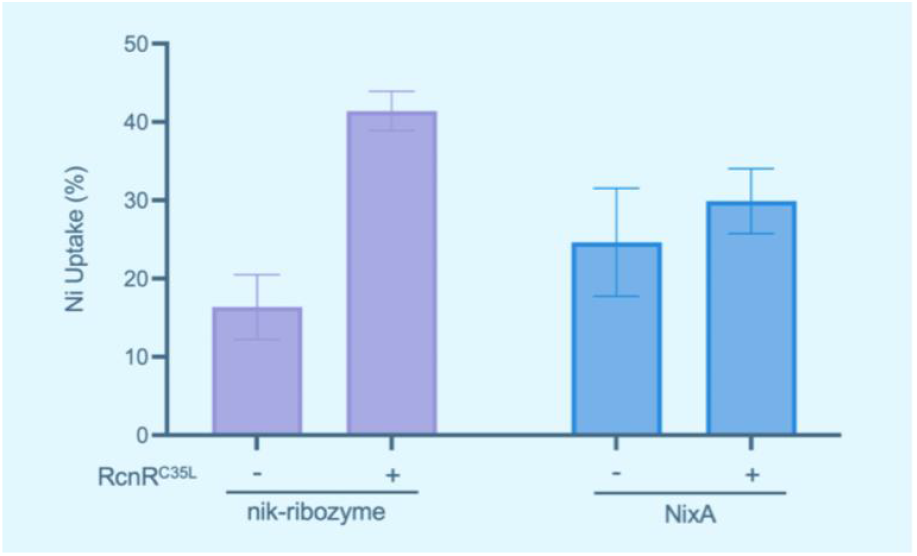
Comparison of Ni^2+^ Uptake Efficiency, with and without RcnR^C35L^. The graph shows the percentage of Ni^2+^ absorbed by *E. coli* expressing different constructs after 5 hours of growth in a medium containing 20 mg/L Ni^2+^ (*E. coli* strain: BL21 DE3, induced with 1 mM IPTG). Ni^2+^ uptake was calculated based on the difference between initial and final concentrations in the supernatant, divided by 20 mg/L. The optical density (OD_600_) of the initial bacterial suspension was adjusted to 0.5. Culture for 5 hours, at 37°C with a rotating speed at 220 rpm. Three biological replicates were performed for each condition, and error bars represent the standard errors of the means (SEM) of these replicates. RcnR^C35L^ refers to a mutation in which cysteine (C) at position 35 in the RcnR protein was substituted with leucine (L). The results indicate that *E. coli* expressing RcnR^C35L^ consistently has higher Ni^2+^ uptake efficiency compared to *E. coli* without RcnR^C35L^ expression.

### Improved Nickel Tolerance and Environmental Robustness

To enable bacterial survival under high-nickel stress, we explored the use of the histidine-rich protein Hpn from H. pylori, known for its nickel-binding and detoxification capacity. Growth assays performed in LB media supplemented with varying concentrations of NiSO_4_ (0.5–2 mM) revealed that strains expressing hpn showed significantly improved growth rates and final optical density compared to control strains (Figure 3). Additionally, nickel accumulation assays confirmed that the protective effect of Hpn indirectly facilitated greater net uptake of nickel by mitigating cytotoxicity (Figure 4).

**Figure 3:**
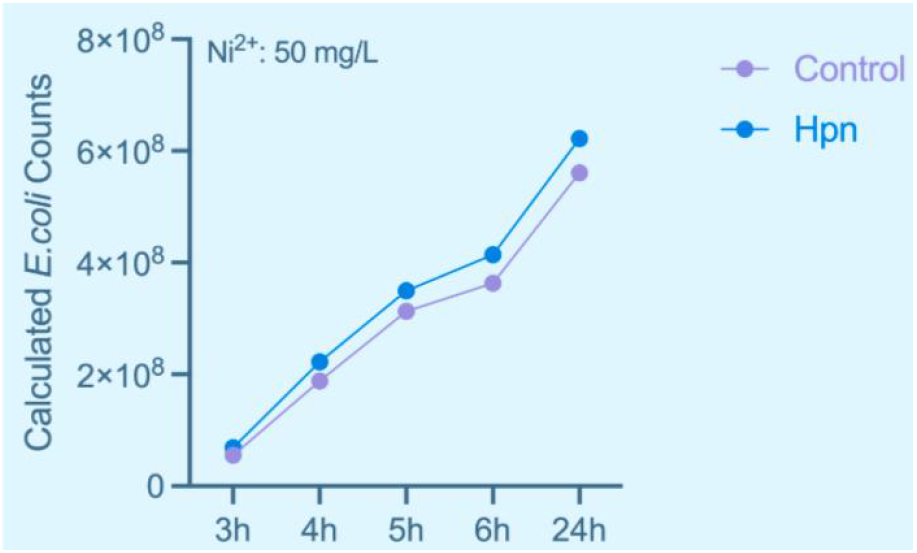
Comparison of *E. coli* Growth curve with and without Hpn in 50 mg/L Ni^2^. The graph illustrates the effect of Ni^2+^ on the growth of *E. coli* expressing Hpn compared to *E. coli* without Hpn expression in a medium containing 50 mg/L Ni^2+^ (*E. coli* strain: BL21 DE3, induced with 1 mM IPTG). The optical density (OD_600_) of the initial bacterial suspension was adjusted to 0.5. *E. coli* growth was measured by OD_600_, and the bacterial counts were calculated using a standard conversion, where OD_600_ = 1 corresponds to 5.39 × 10^8^ cells. The results indicate that *E. coli* expressing Hpn has greater tolerance to Ni^2+^, exhibiting higher growth rates than *E. coli* without Hpn expression under the same conditions.

**Figure 4:**
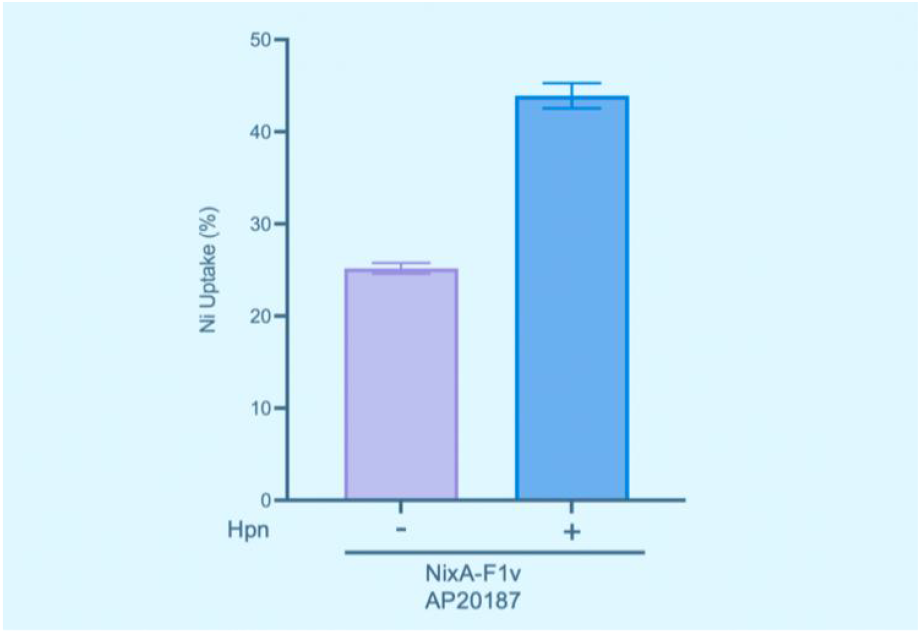
Comparison of Ni^2+^ Uptake Efficiency, with and without Hpn. The graph shows the percentage of Ni^2+^ absorbed by *E. coli* expressing different constructs after 5 hours of growth in a medium containing 50 mg/L Ni^2+^ (*E. coli* strain: BL21 DE3, induced with 1 mM IPTG). Ni^2+^ uptake was calculated based on the difference between initial and final concentrations in the supernatant, divided by 50 mg/L. The optical density (OD_600_) of the initial bacterial suspension was adjusted to 0.5. Culture for 5 hours, at 37°C with a rotating speed at 220 rpm. Our best nickel uptaker NixA-F1v was used, and AP20187 was added to induce NixA dimerization.

Beyond heavy metal stress, we also addressed the vulnerability of engineered strains to bacteriophage infection. Expression of the membrane protein YejM, previously implicated in outer membrane remodeling, conferred enhanced resistance to phage exposure. In phage challenge assays, yejM-expressing *E. coli* maintained significantly higher survival rates relative to controls lacking the gene (Figure 5), indicating the utility of yejM in maintaining strain stability under environmental threats.

**Figure 5:**
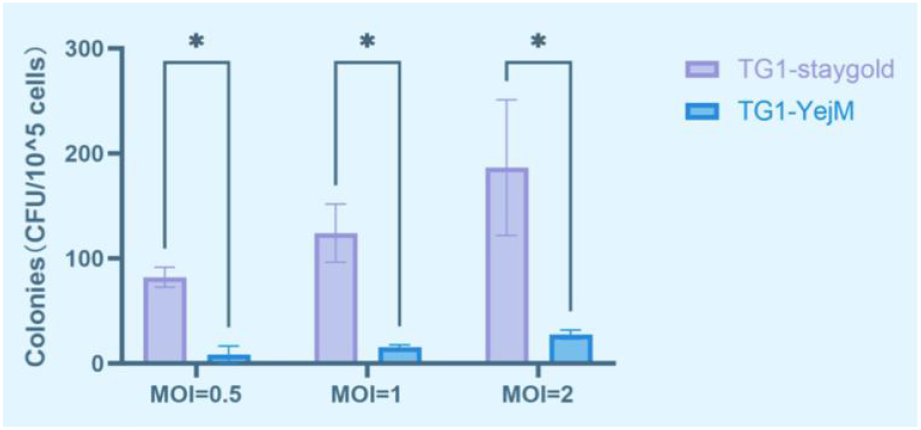
Phage Colony Formation, with and without YejM expression. *E. coli* TG1 carrying either stayGold fluorescent protein and YejM under the J23107 promoter were infected with M13KO7 phage at different MOIs. Colonies were counted after incubating at 37°C for 16 hours, on selection LB plates. “*” indicates a p-value less than 0.01. Under any MOI condition, the colony count of *E. coli* TG1 carrying YejM was significantly lower than that of stayGold, indicating that YejM expression confers resistance to phage infection.

### Carboxysome Targeting via the EP Sequence

To establish a spatially confined environment for enhanced redox catalysis, we aimed to direct functional enzymes such as hydrogenase into the carboxysome by fusing them with the EP (Encapsulation Peptide) targeting signal. As a proof-of-concept, we constructed a fusion protein consisting of the fluorescent marker Staygold and the EP sequence. This design allowed us to visually track subcellular localization in the presence or absence of assembled carboxysomes. The differential localization of Staygold-EP, contingent upon the presence of carboxysome components, validates the feasibility of using the EP signal to direct catalytic proteins into engineered bacterial microcompartments(Figure 6). This strategy lays the foundation for the compartmentalized assembly of hydrogenase in a dedicated reduction chamber, potentially enhancing catalytic efficiency by spatial proximity and substrate channeling.

**Figure 6:**
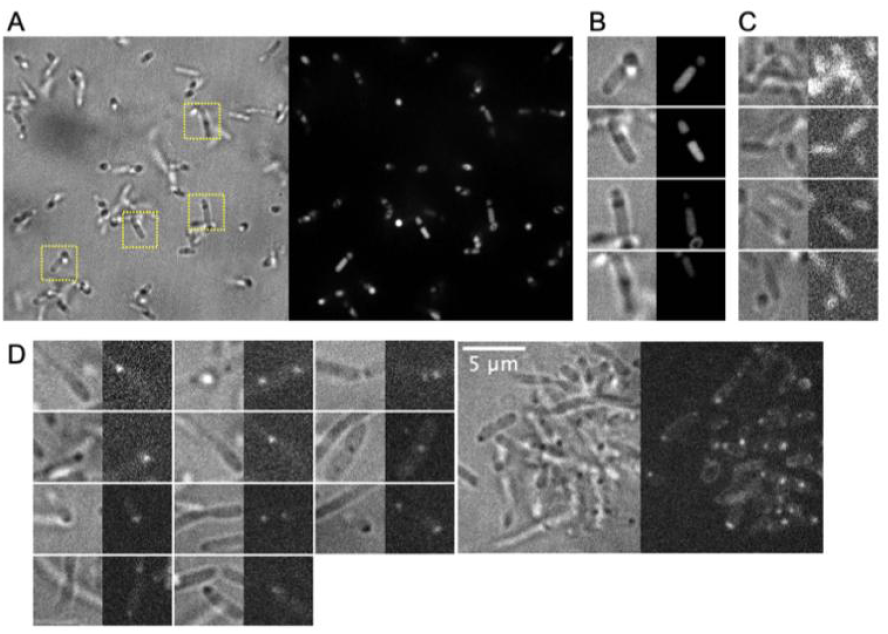
Fluorescence images of *E. coli* expressing Staygold fused with EP, without or with cso-S3. Images were captured using spinning disk confocal with a 150x objective lens. Bacteria in A-C only express stayGold fused with EP, while bacteria in D simultaneously express stayGold fused with EP and cso, without csoS3. 1 mM IPTG was added to A,B only. Images without scale bar are 5×5 µm square, unless specifically indicated below. (A) The entire image field is shown (41.27×41.27 µm square), with brightfield image on the left, and green fluorescence image on the right. (B) Four regions in (A) are enlarged, showing uniform distribution of green fluorescence. (C) Although no IPTG was added, leaky expression from the promoter is sufficient to fill bacteria with green. (D) With all carboxysome components expressed, stayGold fused with EP concentrated to the carboxysome. Leaky expression from the promoter is sufficient to drive 1 or 2 carboxysome formed within each bacteria.

### Intracellular Biomineralization of Nickel Microparticles

To convert accumulated nickel into a reusable form, we introduced a synthetic biomineralization module (F module) into *E. coli*. Upon exposure to 1 mM NiSO_4_, cells expressing the F module exhibited visible particle formation, confirmed via electron microscopy and elemental mapping (Figure 7). The composition and morphology of the particles were consistent with the formation of nickel microparticles, supporting the hypothesis that intracellular biomineralization had occurred.

**Figure 7:**
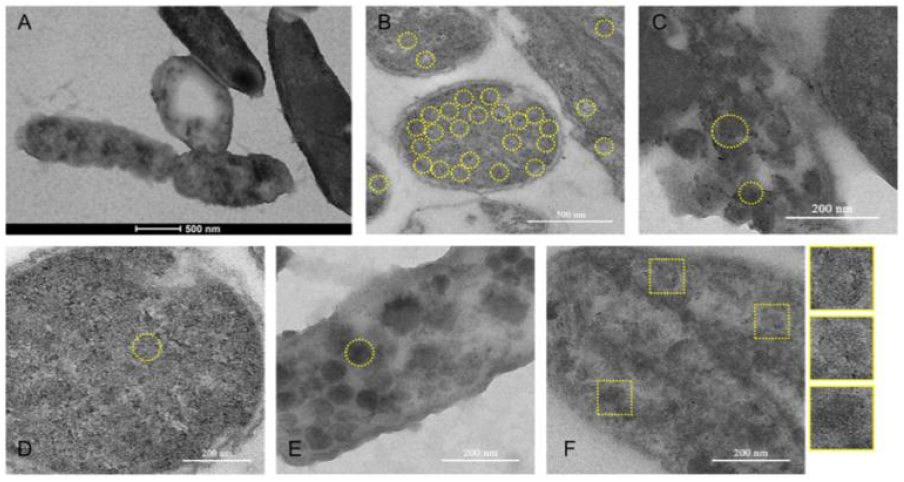
TEM images of negative stained *E. coli* expressing the F module. Osmium tetroxide and uranyl acetate were used for the staining A-E. Scale bar shown on the image. (A) Overview of *E. coli* cells. (B) Sections of bacteria, filled with carboxysome-sized regions (CSR) surrounded by electron dense dots. In one cell, all visual CSR are circled by yellow dash lines. (C) The size of CSR are various, with two examples circled. (D,E) For cells with less electron dense dots, CSR are clear, with the cell in C fully packed with CSR and the cell in D sparsely packed. (F) No uranyl acetate staining. The image confirms that the electron dense dots throughout A-E are not salt crystals but actual metallic particles, which we believe are Ni particles. Three 80-nm square regions are enlarged, showing polyhedral outline of CSR, with Ni particles surrounded.

Unexpectedly, we observed that strains expressing the F module alone—without any engineered nickel uptake system—accumulated significantly more nickel than control strains, and even surpassed strains harboring the U module alone (Figure 8). This finding suggests the possibility of passive uptake or enhanced retention facilitated by the biomineralization pathway, warranting further mechanistic investigation.

**Figure 8:**
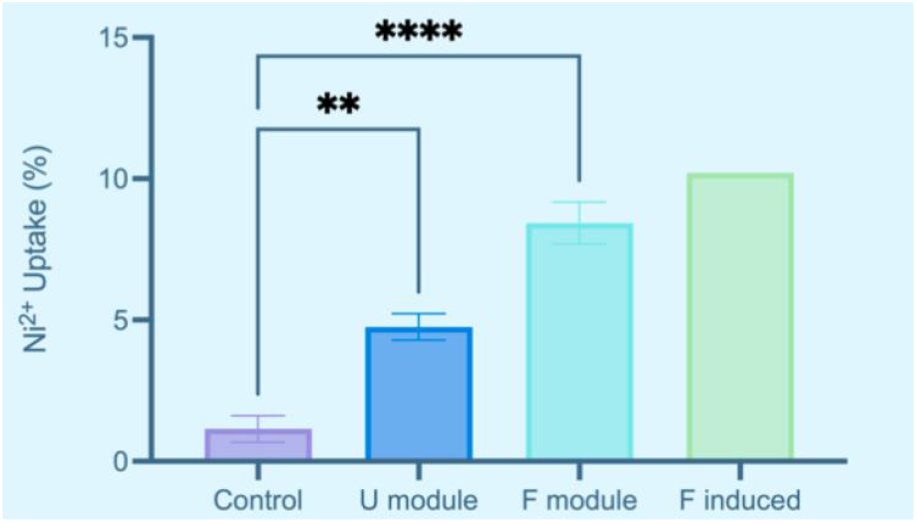
TEM images of negative stained *E. coli* expressing the F module. The graph shows the percentage of Ni^2+^ absorbed by *E. coli* expressing indicated modules (*E. coli* strain: BL21 DE3). Ni^2+^ uptake was calculated based on the difference between initial and final concentrations in the supernatant, divided by 100 mg/L. The single bacteria colony was picked and grown overnight to reach optical density (OD_600_) > 1. Prepare a sealed 25-mL LB culture in a 250-mL bottle, with: 100 µL overnight bacteria liquid culture, 25 µg/mL Kan, 1 mM methyl viologen dichloride, 100 mg/L NiCl_2_, bubbled with ~250 mL 5.6% hydrogen gas (slowly, with hand-shaking, about 5 minutes). Culture for 30 hours, at 37°C with a rotating speed at 220 rpm. Four biological replicates were performed for each condition, and error bars represent the standard errors of the means (SEM) of these replicates. Plain BL21 DE3 was used as control. None of the four bacteria with U module was able to grow during overnight culture if induced with 1 mM IPTG, only 1 F module grow, which was further examined by TEM. Additional 1 mM IPTG was added into the 25-mL culture of “F induced”. ANOVA test shows that all constructs increase Ni^2+^ uptake significantly compared to the control (U module, p = 0.0045; F module, p < 0.0001). P value was calculated using Dunnett’s post-test.

**Figure 9:**
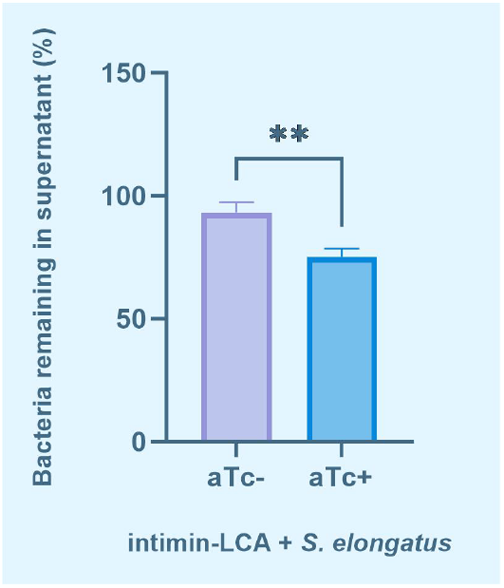
The percentage of bacteria remaining in the supernatant after 6 hours between aTc-induced (+) and uninduced (−) samples. After aTc induction, *E. coli* bacteria express intimin-LCA, which specifically binds with *Synechococcus elongatus*. We used OD_685_ to plot the ratio of the remaining *S. elongatus* in the supernatant to the initial concentration after 6 hours of cultivation. The aTc-induced sample show a significant reduction in the bacteria remaining in the supernatant compared to the uninduced sample, indicating enhanced aggregation between *E. coli* and *S. elongatus*. “**” indicates statistical significance less than 0.01.

### Establishing Hydrogen Supply Through Synthetic Adhesion

To support long-term bioprocesses such as microbial electrochemical systems or co-culture remediation, we designed a synthetic adhesion system to tether *E. coli* to hydrogen-producing cyanobacteria. Surface display elements were introduced to both partners, facilitating stable physical interactions. Microscopy confirmed adhesion between engineered *E. coli* and Synechococcus elongatus under co-culture conditions (Figure 8). Although hydrogen production was not directly quantified in this experiment, the successful establishment of a stable interspecies consortium lays the foundation for coupling hydrogen supply with nickel bioremediation in future systems.

## Methods

### Overlap Extension PCR (OE PCR)

The fragments were amplified separately using standard PCR conditions. The two amplified fragments were then mixed and used as templates in a second round of PCR to extend the overlapping regions and form a full-length product. The PCR products were analyzed by agarose gel electrophoresis, purified using a gel extraction kit, and cloned into an appropriate vector for further applications. The primers included 20-30 base pairs of complementary sequences to ensure efficient extension, and the success of the process was confirmed by sequencing of the PCR products.

### Ni2+ Concentration Measurement

Ni^2+^ concentration was determined by a dimethylglyoxime colorimetric assay. A set of NiCl_2_·6H_2_O standards (0–1.8 mg L^−1^) was prepared in deionised water and treated in parallel with the samples. For each reaction, the required volume of standard or sample was combined with deionised water to 800 µL, followed by 100 µL of 0.05 % (w/v) dimethylglyoxime in ethanol, 90 µL of 1 M ammonium-citrate buffer (pH 9.0) and 10 µL of 5 % (w/v) Na_2_CO_3_ in a 1.5 mL microcentrifuge tube, giving a final volume of 1 mL. The mixtures were vortexed and incubated at 25 °C for 10 min to allow complex formation, then the absorbance at 520 nm was recorded in a UV–visible spectrophotometer using 1 cm path-length quartz cuvettes. A calibration curve was generated by linear regression of absorbance versus nominal Ni^2+^ concentration, and sample concentrations were obtained from the regression equation. Each measurement was performed in triplicate and take average outcome.

### Fluorescence Microscopy Observation

A 1 % (w/v) low-melting agarose solution was prepared in distilled water by microwave heating for 30–60 s until fully dissolved, and 1.0–1.2 mL of the molten agarose was poured onto a glass slide bordered on both long edges by two stacked slides that served as spacers; a second slide was immediately laid on top to create a level surface. After cooling to approximately 40 °C and solidifying for 45–60 min at room temperature, the spacers and top slide were removed, leaving a uniform agarose pad on the base slide. Ten microlitres of cell suspension were applied to the centre of the pad and allowed to settle for 2 min, then a 22 mm × 22 mm coverslip was gently placed to eliminate air gaps. The preparation was examined with an inverted fluorescence microscope fitted with a 100× oil-immersion objective (NA 1.45), LED illumination and filter sets matched to the fluorophores under investigation. Images were acquired with a CMOS camera at 12-bit depth and constant exposure time, and raw data were processed in ImageJ for downstream analysis.

### Transmission Electron Microscopy Sample Preparation and Imaging

Bacterial pellets were fixed in 2.5 % glutaraldehyde in 0.1 M phosphate buffer (pH 7.4) for 2 h at 4 °C, washed six times for 10 min each in the same buffer, post-fixed in 1 % osmium tetroxide prepared in phosphate buffer for 1 h on a clinical rotator, and washed again six times in buffer to remove residual OsO4. Specimens were dehydrated through a graded ethanol series (30 %, 50 %, 70 %, 90 % and 100 %, 10 min each) followed by two changes of 100 % acetone for 10 min. Samples were infiltrated at room temperature with Durcupan ACM resin by sequential incubation in acetone : resin mixtures of 2 : 1, 1 : 1 and 1 : 2 (30 min each) and finally in 100 % resin for 2 h, then embedded in gelatin capsules filled with Durcupan ACM Mixture #2 and polymerised at 60 °C for 48 h. Ultrathin sections (70 nm) were cut with a diamond knife on an ultramicrotome, collected on 200-mesh copper grids, contrasted for 10 min in 2 % aqueous uranyl acetate followed by 5 min in Reynolds lead citrate, air-dried, and examined with a transmission electron microscope operating at 80 kV. Digital micrographs were captured with a CCD camera at native resolution and exported as TIFF files for analysis.

### Cyanobacteria–E. coli Aggregation Assay

Exponentially growing cultures of *Synechococcus elongatus* PCC 7942 and *Escherichia coli* BL21(DE3) were harvested at OD_750_ ≈ 0.6 and OD_600_ ≈ 0.8, respectively, washed twice in BG-11 medium and resuspended to OD_750_ = 0.4 and OD_600_ = 0.5. Equal volumes of the two suspensions were combined in 15 mL glass tubes to a final volume of 6 mL, gently inverted three times, and incubated under continuous illumination (50 µmol photons m^-2^ s^-1^, 30 °C) without shaking. At 0, 30, 60, 120 and 240 min, 1 mL aliquots were withdrawn without disturbing visible flocs, transferred to 1.5 mL microcentrifuge tubes and centrifuged at 1000 × g for 2 min to pellet aggregates; the absorbance of the supernatant was recorded at 750 nm and 600 nm to monitor the residual cyanobacterial and bacterial cell densities. Aggregation efficiency was calculated as (A0 − At)/A0, where A0 and At denote the initial and time-point absorbance values. All measurements were performed in triplicate, and blanks containing individual monocultures were processed in parallel to correct for sedimentation.

## Reference

[1] Rizwan, M., Usman, K., & Alsafran, M. (2024). Ecological impacts and potential hazards of nickel on soil microbes, plants, and human health. Chemosphere, 357, 142028. 10.1016/j.chemosphere.2024.142028

[2] Wang, Z., Yeung, K. W. Y., Zhou, G. J., Yung, M. M. N., Schlekat, C. E., Garman, E. R., Gissi, F., Stauber, J. L., Middleton, E. T., Lin Wang, Y. Y., & Leung, K. M. Y. (2020). Acute and chronic toxicity of nickel on freshwater and marine tropical aquatic organisms. Ecotoxicology and environmental safety, 206, 111373. 10.1016/j.ecoenv.2020.111373

[3] Wang, Z., Kwok, K. W., & Leung, K. M. (2019). Comparison of temperate and tropical freshwater species’ acute sensitivities to chemicals: An update. Integrated environmental assessment and management, 15(3), 352–363. 10.1002/ieam.4122

[4] Global Critical Minerals Outlook 2024 – Analysis. (2024, May 17). IEA. https://www.iea.org/reports/global-critical-minerals-outlook-2024

[5] Diep, P., Leo Shen, H., Wiesner, J. A., Mykytczuk, N., Papangelakis, V., Yakunin, A. F., & Mahadevan, R. (2023). Engineered nickel bioaccumulation in Escherichia coli by NikABCDE transporter and metallothionein overexpression. Engineering in Life Sciences, 23(7), 2200133. 10.1002/elsc.202200133

[6] Mobley, H. L., Garner, R. M., & Bauerfeind, P. (1995). Helicobacter pylori nickel-transport gene nixA: synthesis of catalytically active urease in Escherichia coli independent of growth conditions. Molecular microbiology, 16(1), 97–109. 10.1111/j.1365-2958.1995.tb02395.x

[7] Higgins, K. A., Chivers, P. T., & Maroney, M. J. (2012). Role of the N-terminus in Determining Metal-Specific Responses in the E. coli Ni- and Co-Responsive Metalloregulator, RcnR. Journal of the American Chemical Society, 134(16), 7081–7093. 10.1021/ja300834b

[8] Saylor, Z., & Maier, R. (2018). Helicobacter pylori nickel storage proteins: recognition and modulation of diverse metabolic targets. Microbiology (Reading, England), 164(8), 1059–1068. 10.1099/mic.0.000680

[9] Tang, H., & Hall, M. B. (2017). Biomimetics of NiFe-Hydrogenase: Nickel- or Iron-Centered Proton Reduction Catalysis? Journal of the American Chemical Society, 139(49), 18065–18070. 10.1021/jacs.7b10425

